# Nucleotide-derived bacterial alarmones attenuate the induction of type-I interferon responses in a murine macrophage reporter cell line

**DOI:** 10.1101/2025.01.27.635077

**Authors:** Ryan P Kilduff, Erin B Purcell, Lisa M Shollenberger

**Affiliations:** Department of Biological Sciences at Old Dominion University, Norfolk, VA; Department of Chemistry and Biochemistry at Old Dominion University, Norfolk, VA

## Abstract

The stringent response is a well-studied phenomenon in many bacterial systems and regulates resource-consuming activities such as transcription, translation, and replication. The stringent response is a well-conserved signaling framework, as are the nucleotide-derived signaling mediators, collectively referred to as (p)ppGpp or as alarmones. There is a wealth of research evaluating nucleotide-derived alarmone signaling in bacterial models, however, their potential to modulate innate immune signaling has not yet been evaluated. Several common pathogen-synthesized molecules, such as lipopolysaccharide (LPS) and cyclic-di-AMP (c-di-AMP), act as pathogen-associated molecular patterns (PAMPs), which are common patterns that alert the innate immune system of bacterial infection. The goal of this work is to elucidate the impact of (p)ppGpp on innate immune signaling. To explore this, RAW-Dual cells were incubated with guanosine tetraphosphate (ppGpp) and guanosine pentaphosphate (pppGpp), two well-studied nucleotide-derived alarmones found in many different pathogenic bacteria, as well as with GTP. Both ppGpp and pppGpp were able to significantly reduce the expression of secreted luciferase in RAW-Dual cells in a dose-dependent manner, indicating a reduction of the interferon-stimulated regulatory elements (ISREs). Neither alarmone impacted secreted embryonic alkaline phosphatase (SEAP) secretion, which reports for NF-kB activation. This is the first work to suggest that nucleotide-derived alarmones produced by bacteria may impact an arm of innate immunity responsible for type-I interferon secretion.

**Author Summary:** Many different disease-causing bacteria use a similar signaling network to help them survive in environments with low resources; this network is called “the stringent response.” All bacteria that use this network also produce signaling molecules, called alarmones, that help coordinate their response to resource deprivation. Interestingly, the human immune system recognizes molecules that are made by many pathogens. As such, our team decided to evaluate whether these bacterially-produced alarmones are able to affect our immune system. We discovered, rather than stimulating our immune system, that high concentrations of these alarmone compounds are able to turn off an important arm of the immune system that is essential in combating viral infections. This suggests another mechanism that bacteria may use to hide from the immune system during infection. In addition, there are potential clinical uses for a molecule that is selectively turn off arms of the immune system, and these molecules may have potential as future anti-inflammatory drugs after further research is able to explore their mechanism of action.

## Introduction

The stringent response is a well-studied phenomenon in many bacterial systems and regulates resource-consuming activities such as transcription, translation, and replication (1–4). The stringent response is perhaps best understood through the model organism *Escherichia coli*, in which the presence of uncharged tRNA in the ribosome triggers an increase in the cytosolic concentration of secondary messengers called alarmones (5,6). Guanosine tetraphosphate and guanosine pentaphosphate, collectively referred to as (p)ppGpp are GTP-derived alarmones that mediate the stringent response. Their synthesis and breakdown are primarily dependent on the RelA/SpoT homology (RSH) family of proteins, which contain dual (p)ppGpp synthetase hydrolase domains, such as RelA in *E. coli* (4). Increased cytosolic (p)ppGpp results in transcriptional changes in *E. coli* that slow growth rate and alter metabolism to preserve energy and nutrients, typically resulting in stationary growth (1,3,6), and also promote recovery after growth disruption (7).

The signaling framework of the stringent response has been observed in most bacterial systems evaluated thus far (1,4); however, the mechanism of impact, as well as the effects of (p)ppGpp signaling, can differ greatly between genera. Induction of the stringent response in *E. coli* results in changes to the affinity of RNA polymerase for σ factors such that σS and σ32 outcompete σ70, resulting in an altered transcriptional profile (8); however, ppGpp does not interact with RNA polymerase in Firmicutes, and instead directly activate proteins involved in the stress response (9). In addition to impacting resource conservation and growth rate, induction of the stringent response has also been shown to promote characteristics of virulence such as biofilm formation (9–15), antibiotic resistance (9,10,13–17), toxin production (1,5,11,18,19), and spore formation (15,19,20) in several human pathogens.

In addition to (p)ppGpp, several bacteria are known to produce another class of nucleotide-derived messengers: cyclic dinucleotides. Among these messengers are c-di-AMP and c-di-GMP, which are synthesized from two ATP or GTP molecules, respectively, and are produced by pathogens such as *Listeria monocytogenes* (21–24) and *Clostridoides difficile* (25–27). Interestingly, *L. monocytogenes* secretes c-di-AMP through the Mdr family of multi-drug efflux pumps, as prolonged, elevated intracellular levels of this messenger can negatively impact the expression of virulence factors and can be toxic to the organism (21,28). This makes c-di-AMP available for recognition by STING, an ER-resident innate immune receptor, which responds to the recognition of secreted cyclic dinucleotides by stimulating the release of type-I interferons through IRF3 (22,29,30).

Cyclic dinucleotides, such as c-di-AMP and c-di-GMP, are just one class of pathogen-associated molecular patterns (PAMPs) known to stimulate the immune system through pattern recognition receptors (PRRs), such as STING (29,31). PRR recognition of PAMPs is a vital step in stimulating the innate immune system and is essential for subsequently activating the adaptive immune system (32,33). An “ideal” PAMP is a molecular pattern that is found in a wide range of organisms, such as flagellin, lipopolysaccharide (LPS), and CpG motifs in DNA, all of which are recognized in humans by a class of PRRs called toll-like receptors (TLRs), resulting in the induction of signaling through NF-κB (32). Despite the wealth of research evaluating (p)ppGpp signaling in bacterial models, their potential to modulate innate immune signaling has not yet been evaluated. Considering the ubiquity of (p)ppGpp signaling in bacteria, and also considering the ability of host receptors to recognize nucleotide-derived bacterial messengers (e.g. cyclic dinucleotides through STING), alarmones released by pathogenic bacteria may play a role in the stimulation (or attenuation) of innate immune signaling.

## Methods

### Cell culture

The RAW-Dual cell line (InvivoGen), a dual reporter mouse macrophage line, was cultured according to manufacturer specifications and with institutional biosafety approval (24-010). To evaluate immune responses, cells were seeded in a 96-well plate at 2.0 x 10^5^ cells per well and incubated with alarmone (ppGpp or pppGpp, Jena Bioscience), c-di-AMP (InvivoGen), LPS O111:B4 (Sigma), GTP (Biobasic), and/or Dexamethasone (Sigma) for 23h at 37°C and 5% CO_2_ in DMEM media without selective antibiotics. After incubation, the luciferase and SEAP assays were completed according to manufacturer instructions (InvivoGen).

### Analysis

Data were normalized within experiment to the mean of media-only controls (single incubations) or to the mean of of LPS- or c-di-AMP-treated controls (co-incubations), then pooled from 12 independent experiments. Significant differences across means of individual treatment groups were compared using the Kruskal-Wallis test, and means between individual groups were compared using Dunn’s multiple comparison test. Three outliers were identified and removed using the Grubb’s test. Differences between treatments were analyzed by comparing the area under the curve (AUC) over 0-1000uM concentrations of all treatments, and differences between the AUC between treatments were analyzed by Brown-Forsythe ANOVA. All graphical figures and statistical analyses were conducted within GraphPad Prism version 10.2.3. Normalized data is available on Mendeley Data <doi: 10.17632/d64fcb5jyc.2>.

## Results

RAW-Dual cells were first incubated with serial concentrations of ppGpp, pppGpp, GTP, LPS, c-di-AMP or dexamethasone. LPS at a concentration of 1ug/mL significantly stimulated secretion of both luciferase and SEAP, whereas c-di-AMP at 20uM increased the secretion of Luciferase compared to media, but not SEAP (**Figure S1**). Interestingly, luciferase expression shows a trend of attenuation below baseline levels with increasing concentrations of both ppGpp and pppGpp (**Figure 1A**), although this same phenomenon is not seen with SEAP secretion (**Figure 1B**). To prove this phenomenon is not a reaction between (p)ppGpp and Lucia luciferase, supernatants from LPS-incubated RAW-Dual cells, containing high quantities of secreted Lucia luciferase, were mixed with ppGpp, pppGpp and GTP to the highest tested concentrations of 1000uM for (p)ppGpp and 2000uM for GTP. No significant difference in luminescence was observed between any groups containing supernatants exposed to any of the treatments (**Figure S2**), suggesting the attenuated secretion of luciferase in (p)ppGpp-treated cells is due to a cell signaling phenomenon rather than direct interactions with the secreted Lucia luciferase. Differences between the pattern of luciferase signaling between 0uM (media-only) and 1000uM of each treatment were evaluated by AUC, suggesting that the impact of ppGpp (but not pppGpp) on luciferase signaling over that range is distinct from that of GTP (**Figure 1C**).

**Figure 1.**
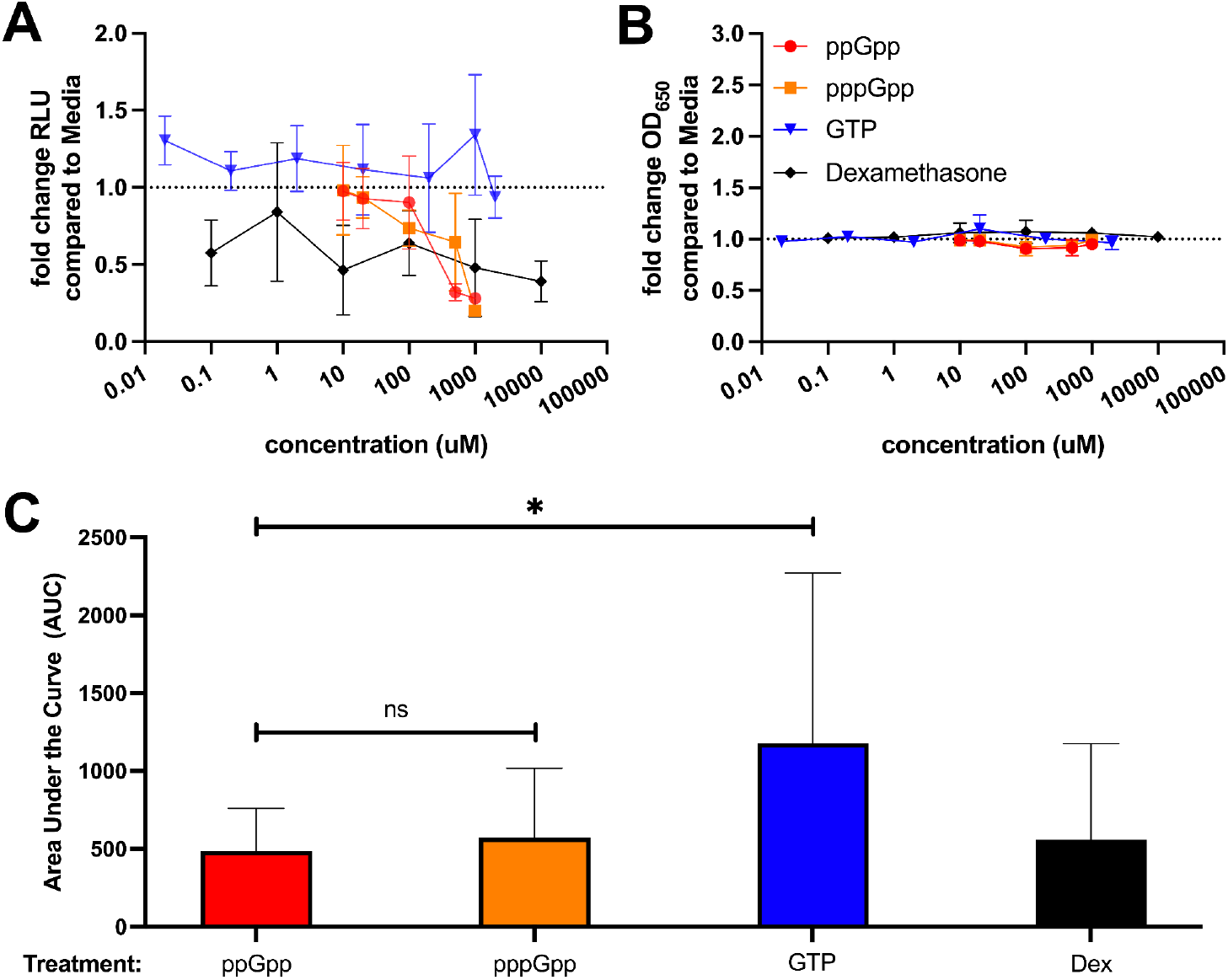
Relative expression of (**A**) RLU in the secreted luciferase assay or (**B**) OD650 in the SEAP assay from supernatants of RAW-Dual cells incubated with serial concentrations of ppGpp, pppGpp, GTP or dexamethasone. **(C)** AUC values for all treatments between 0-1000uM for the secreted luciferase assay; significance is evaluated using Brown-Forsythe ANOVA.

The downward trend of luciferase signaling with increasing concentrations of (p)ppGpp in **Figure 1A** suggests a removal of baseline luciferase signaling. While interesting, we were curious to determine if these compounds could also attenuate luciferase secretion in cells that were treated with known immunostimulatory molecules, such as LPS and c-di-AMP. RAW-dual cells were incubated in media supplemented with 1ug/mL of LPS (**Figure 2**) or 20uM of c-di-AMP (**Figure 3**), in addition to serial concentrations of the compounds ppGpp, pppGpp, and GTP, and dexamethasone. There is a significant, concentration-dependent reduction in luciferase signaling of LPS-incubated RAW-Dual cell supernatants between the doses of 20uM and 500uM of ppGpp and pppGpp (**Figure 2A**). Interestingly, this change is not seen in GTP, despite a chemical similarity to ppGpp and pppGpp. The negative control, dexamethasone, appears to diminish luciferase expression both basally and in LPS-induced RAW-Dual cells (**Figures 1A and 2A**). Interestingly, dexamethasone appears to leave basal SEAP unaffected when not otherwise stimulated (**Figure 1B**), but elevates SEAP expression in lower doses when co-administered with LPS (**Figure 2B**). A Browne-Forsythe ANOVA on the AUC of luciferase over 0uM to 1000uM indicates that both alarmone compounds exhibit an independent effect on luciferase signaling compared to GTP, while ppGpp (but not pppGpp) exhibits an effect independent of dexamethasone (**Figure 2C**). Co-incubation of RAW-Duals with either alarmone or GTP does not appear to exhibit a concentration-dependent impact on luciferase nor SEAP expression (**Figure 3A and 3B**). The effect of either alarmone on luciferase signaling in RAW-Duals over the concentrations of 0uM-1000uM is not significantly different than GTP (**Figure 3C**).

**Figure 2.**
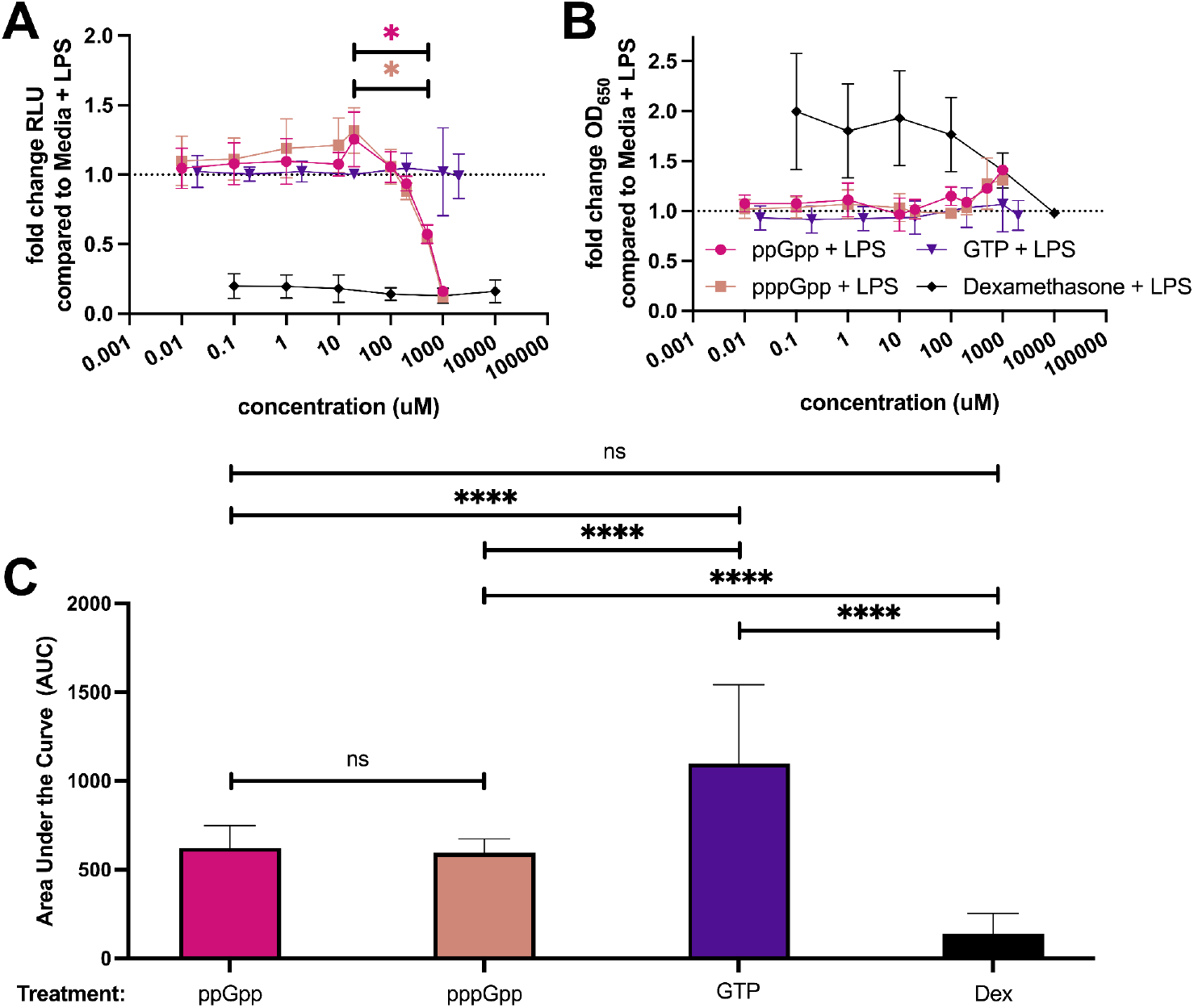
Relative expression of (**A**) RLU in the secreted luciferase assay or (**B**) OD650 in the SEAP assay from supernatants of RAW-Dual cells incubated in 1ug/mL of LPS 0111:B4, in addition to serial concentrations of ppGpp, pppGpp, GTP or dexamethasone. Significance is evaluated using Kruskal-Wallis ANOVA, and p-values were compared using Dunn’s Multiple Comparisons test. **(C)** AUC values for all treatments between 0-1000uM for the luciferase assay; significance is evaluated using Brown-Forsythe ANOVA.

**Figure 3.**
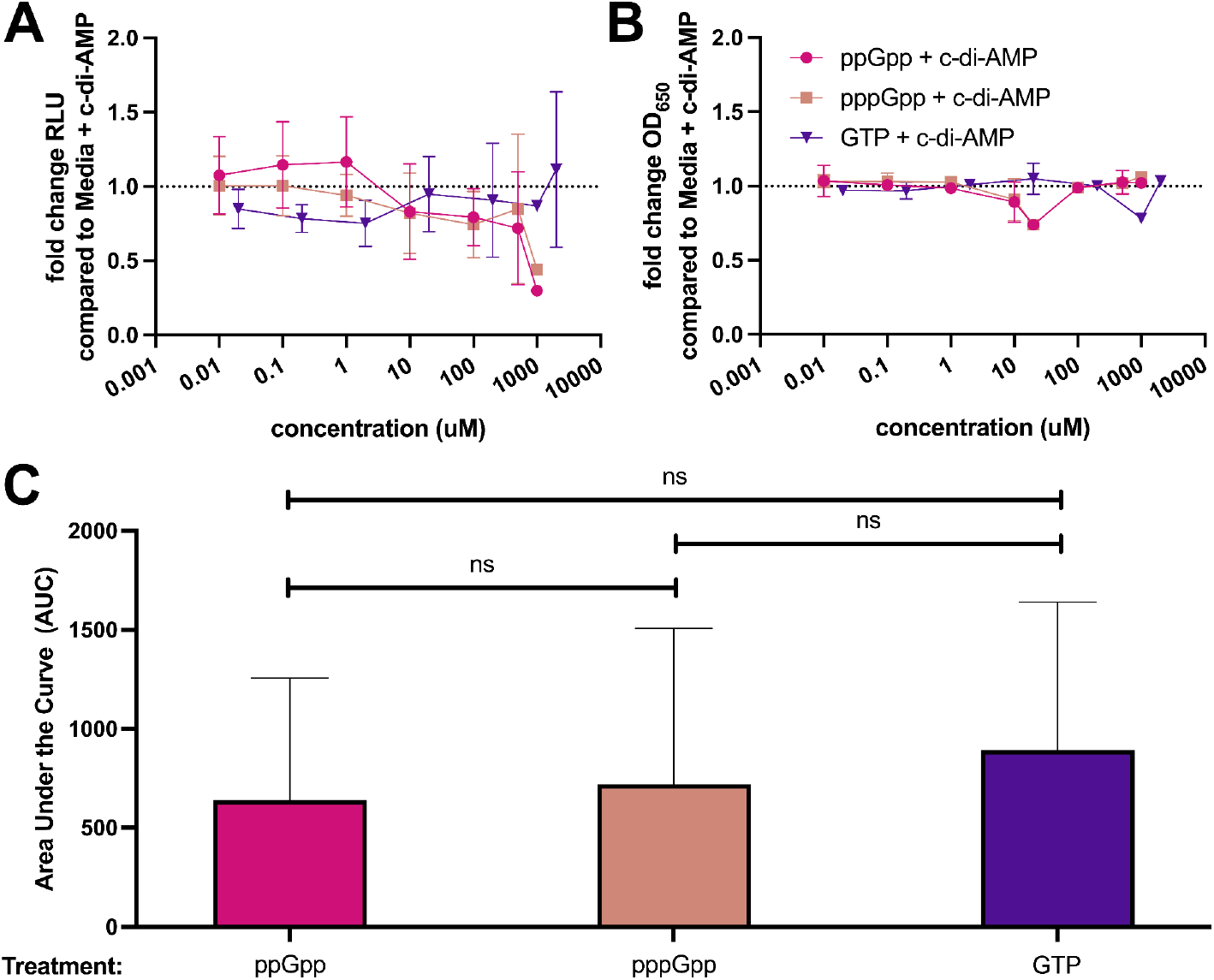
Relative expression of (**A**) RLU in the secreted luciferase assay or (**B**) OD650 in the SEAP assay from supernatants of RAW-Dual cells incubated in 20uM of c-di-AMP, in addition to serial concentrations of ppGpp, pppGpp, or GTP. **(C)** AUC values for all treatments between 0-1000uM for the luciferase assay; significance is evaluated using Brown-Forsythe ANOVA.

**Figure 4.**
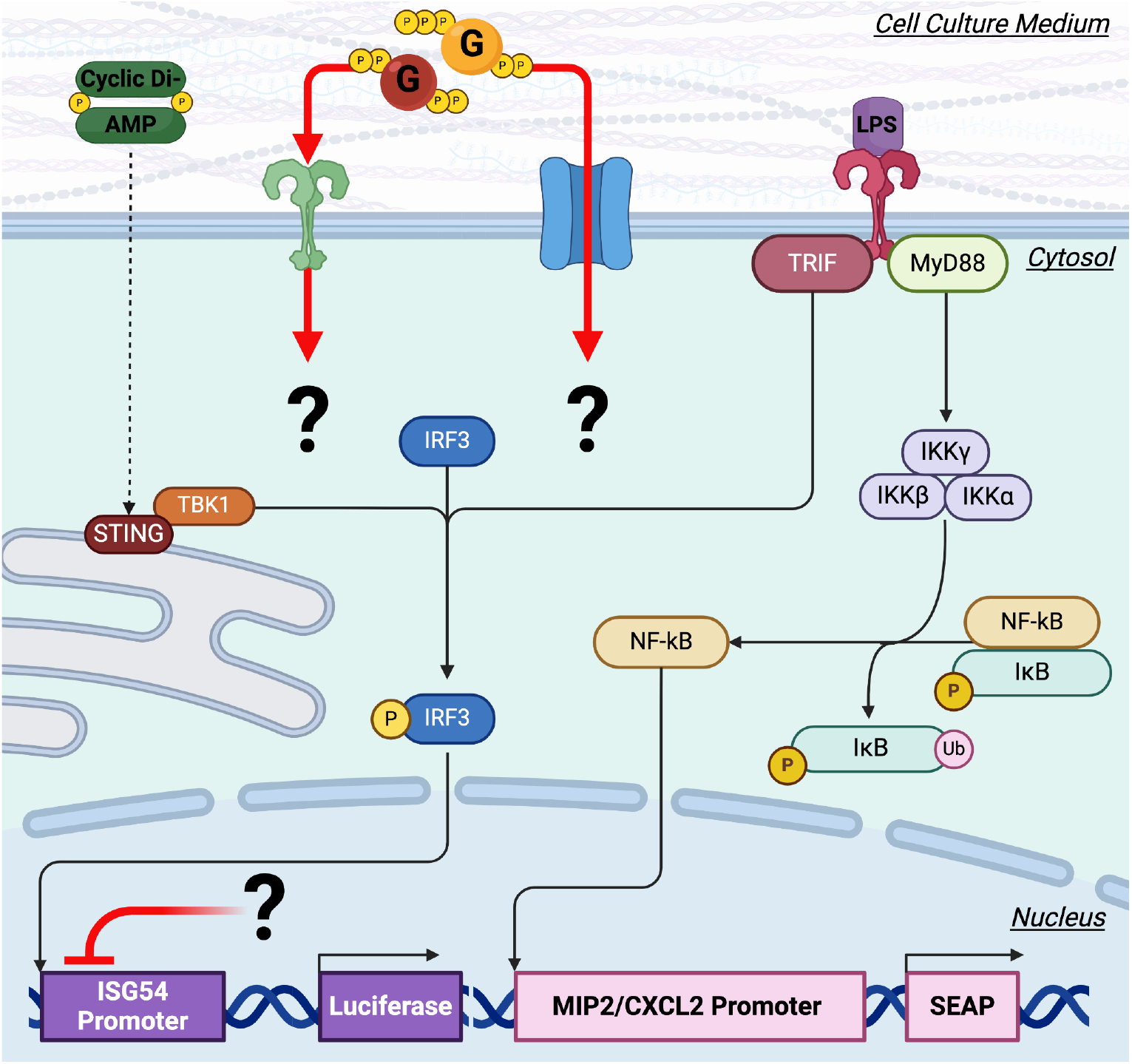
Graphical abstract. Signaling in RAW-Dual cells exposed to c-di-AMP, LPS, and the alarmone compounds, including gaps in the mechanism. It is misunderstood whether the alarmone compounds are able to cross the cell membrane, or if they signal through a currently unidentified receptor. Any potential signaling intermediates are also not currently understood.

Incubating RAW-Dual cells with ppGpp, pppGpp, GTP, and c-di-AMP at any concentration was insufficient to significantly impact production of SEAP when compared to media-only controls (**Figures 1B, 2B, 3B**). When co-incubated with 1ug/mL or 20uM of c-di-AMP of LPS, none of the tested concentrations of ppGpp, pppGpp, nor GTP exhibited dose-dependent impacts on SEAP signaling. This also suggests that the reduction in luciferase secretion of (p)ppGpp-incubated RAW-Dual cells is likely a signaling phenomenon; If (p)ppGpp were simply toxic to cells, then it is likely that values for both the luciferase and SEAP assays would both be attenuated with increasing concentrations when compared to media controls.

## Discussion

These findings suggest that GTP-derived alarmones (p)ppGpp may influence innate immune signaling in murine macrophages through the attenuation of signaling through ISREs, but not NF-kB. Both ppGpp and pppGpp attenuated luciferase secretion in RAW-Duals by themselves (**Figure 1A**) and significantly attenuated luciferase secretion of RAW-Duals co-incubated with LPS in a dose-dependent manner (**Figure 2A**) without impacting SEAP secretion (**Figure 2B**). There was no significant reduction of c-di-AMP induced luciferase secretion in RAW-Dual cells incubated at higher concentrations of either alarmone compound. There may be several reasons for this. Firstly, c-di-AMP is known to stimulate the same pathway that (p)ppGpp seems to attenuate, so the specific mechanism of signaling may be more complicated than a simple linear inhibition pattern. In addition, c-di-AMP is not as potent an inducer of luciferase secretion in RAW-Dual cells compared to LPS (**Figure S1**), so the differences may simply be on a different scale that may be elucidated through further work. As luciferase and SEAP secretion report for ISRE and NF-KB activation, respectively, the findings in this manuscript suggest that the alarmones (p)ppGpp are able to modulate ISRE-based signaling while leaving NF-KB-based signaling unaffected.

Dexamethasone was used in this work as a negative control, as it is known to reduce signaling through NF-KB (34). Interestingly, however, our work showed that dexamethasone did reduce impact basal expression of SEAP at any tested concentration, and actually appeared to elevate SEAP secretion at lower concentrations. One explanation for this may involve the kinetics of the response, as previous work has described that administration of dexamethasone before infection resulted in stronger reduction of TLR-induced NF-KB signaling compared to treatment during active infection (35). As this experiment resulted in co-treatment of dexamethasone and LPS, rather than pre-treatment with dexamethasone, this may explain the elevated expression of NF-KB at lower levels, although this pattern should be further investigated. Despite this, the alarmone compounds do not appear to impact SEAP secretion over the tested ranges, suggesting the impact on signaling between alarmones and dexamethasone is distinct.

Cyclic dinucleotides, such as c-di-AMP secreted by *Listeria monocytogenes*, are secreted during infection and are known to stimulate the innate immune system (21). Here, we show that alarmones, which are reciprocally upregulated with cyclic dinucleotides (36–39), downregulate the inflammatory responses to cyclic dinucleotides, providing a possible mechanism for why cyclic dinucleotide expression during active infection is not subject to negative selective pressure.

