## Supplemental figures for "Nucleotide-derived bacterial alarmones attenuate the induction of type-I interferon responses in a murine macrophage reporter cell line"

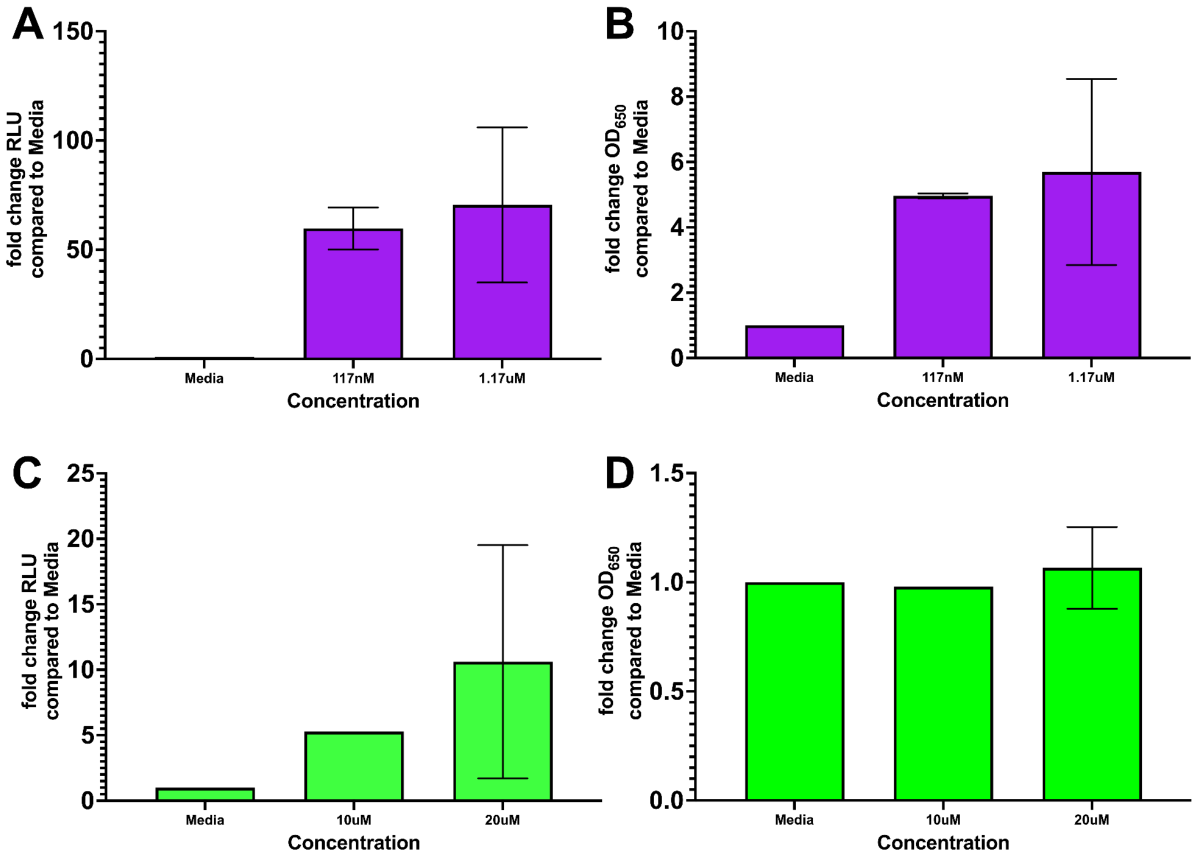


**Figure S1.** Relative expression of secreted Lucia luciferase (**A, C**) or secreted embryonic alkaline phosphatase (**B, D**) of RAW-Dual cells incubated with several concentrations of LPS (**A, B)** or c-di-AMP (**C, D**). Values for each individual replicate were normalized to values from media without any treatments, and data is expressed as the mean fold change in RLU (**A, C**) or OD650 (**B, D**) of cell culture supernatants.

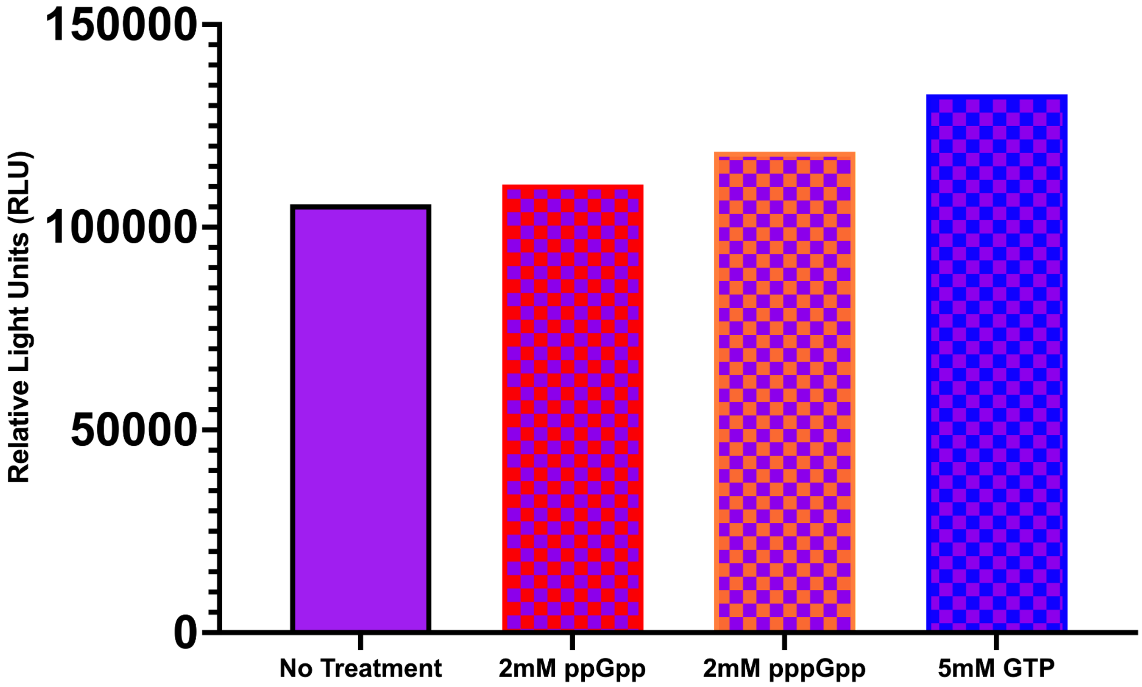


**Figure S2.** Luminescence in relative light units (RLU) of supernatants from RAW-Dual cells incubated in 1ug/mL of LPS 0111:B4 alone, as well as in combination with the treatments ppGpp, pppGpp, GTP. The data shown is a representative example of 3 replicates, in which neither alarmone nor GTP reduced the luminescence in Lucia luciferase-containing supernatants of LPS-incubated RAW-Dual cells.
